# Sex-specific prefrontal-hypothalamic control of behavior and stress responding

**DOI:** 10.1101/2023.07.09.548297

**Authors:** Derek Schaeuble, Tyler Wallace, Sebastian A. Pace, Shane T. Hentges, Brent Myers

**Affiliations:** Biomedical Sciences, Colorado State University, Fort Collins, CO, USA; Integrative Physiology and Neuroscience, Washington State University, Pullman, WA, USA

**Keywords:** corticosterone, glucose, place preference, posterior hypothalamus, social motivation, ventromedial prefrontal cortex

## Abstract

Depression and cardiovascular disease are both augmented by daily life stress. Yet, the biological mechanisms that translate psychological stress into affective and physiological outcomes are unknown. Previously, we demonstrated that stimulation of the ventromedial prefrontal cortex (vmPFC) has sexually divergent outcomes on behavior and physiology. Importantly, the vmPFC does not innervate the brain regions that initiate autonomic or neuroendocrine stress responses; thus, we hypothesized that intermediate synapses integrate cortical information to regulate stress responding. The posterior hypothalamus (PH) directly innervates stress-effector regions and receives substantial innervation from the vmPFC. In the current studies, circuit-specific approaches examined whether vmPFC synapses in the PH coordinate stress responding. Here we tested the effects of optogenetic vmPFC-PH circuit stimulation in male and female rats on social and motivational behaviors as well as physiological stress responses. Additionally, an intersectional genetic approach was used to knock down synaptobrevin in PH-projecting vmPFC neurons. Our collective results indicate that male vmPFC-PH circuitry promotes positive motivational valence and is both sufficient and necessary to reduce sympathetic-mediated stress responses. In females, the vmPFC-PH circuit does not affect social or preference behaviors but is sufficient and necessary to elevate neuroendocrine stress responses. Altogether, these data suggest cortical regulation of stress reactivity and behavior is mediated, in part, by projections to the hypothalamus that function in a sex-specific manner.

## 1. Introduction

Repeated or prolonged stress exposure burdens mental and physical health (de Kloet et al., 2005), predisposing individuals to major depressive disorder (MDD) and cardiovascular disease (CVD) (Ginty et al., 2013; Johnson and Grippo, 2006). Although the comorbidity of MDD and CVD is well-documented, the biological underpinnings remain undetermined. Additionally, the female prevalence of comorbid MDD and CVD is more than double that of males (Goldstein et al., 2019; Möller-Leimkühler, 2007; Rich-Edwards et al., 2018; Wake and Yoshiyama, 2009). These findings collectively highlight the importance of studying stress as a risk factor for mood and cardiometabolic dysfunction, as well as the need to investigate biological sex as a variable (Rich-Edwards et al., 2018). To determine how stressors engage sex-specific neural circuitry and integrate affective and physiological systems, the current report examines prefrontal-hypothalamic circuit regulation of social and motivational behaviors as well as endocrine and autonomic physiology.

The human ventromedial prefrontal cortex (vmPFC) processes emotional and social stimuli (Beckmann et al., 2009; Vijayakumar et al., 2017); moreover, clinical neuroimaging associates the vmPFC with MDD and related symptoms (Drevets et al., 1997; Mayberg et al., 1999). The putative homolog of the vmPFC in rodents, the infralimbic cortex (IL), is also implicated in stressor appraisal and depression-related behaviors (Covington et al., 2010; Hamani et al., 2012; McKlveen et al., 2015; Wallace and Myers, 2023). The vmPFC is composed primarily of glutamatergic pyramidal neurons with output regulated by GABAergic interneurons (Kubota, 2014; McKlveen et al., 2019; Page and Coutellier, 2019). In male rats, glutamate output from the vmPFC regulates coping behaviors (Pace et al., 2020) and is necessary to limit neuroendocrine and autonomic stress reactivity (Myers, 2017; Schaeuble et al., 2019). Also, stimulation of male vmPFC pyramidal neurons increases social motivation and reduces stress responding; however, female vmPFC pyramidal neuron stimulation does not alter social or motivational behaviors and increases stress-induced glucose mobilization and tachycardia (Wallace et al., 2021). Importantly, the vmPFC does not directly innervate the cells that initiate sympathetic or neuroendocrine responses, suggesting the need for intervening circuitry. Quantification of vmPFC projections throughout the forebrain identified the posterior hypothalamus (PH) as a major target (Wood et al., 2019). Additionally, the PH provides glutamatergic innervation of neuroendocrine and pre-sympathetic cell groups (Myers et al., 2016; Nyhuis et al., 2016; Ulrich-Lai et al., 2011; Vertes and Crane, 1996), suggesting a role in stress integration (Schaeuble and Myers, 2022). Functionally, the male PH facilitates HPA axis and cardiovascular stress responses, as well as avoidance behaviors (DiMicco and Abshire, 1987; Myers et al., 2016; Nyhuis et al., 2016). Thus, the PH may serve as an integrator of cortical information to trans-synaptically regulate physiological and behavioral adaptation.

To determine whether vmPFC innervation of the PH regulates behavior or stress responding, we employed circuit-specific approaches to stimulate or inhibit vmPFC synapses in the PH. Stimulation of the vmPFC-PH circuit was carried out via optogenetics to examine whether the circuit mediated affective behaviors or stress responding in male and female rats. Additionally, an intersectional genetic approach relying on tetanus toxin (TeNT) was used to inhibit PH-projecting vmPFC neurons and test the necessity of the vmPFC-PH circuit for stress responding in both sexes. Ultimately, these data provide evidence that the vmPFC-PH circuit sex-specifically integrates behavior and physiology.

## 2. Methods

### 2.1 Experimental design

The current report contains data from 2 separate experiments, each composed of distinct male and female cohorts. A timeline for experiment 1 is outlined (**Fig. 1A**). Experiment 1 investigated n = 17 yellow fluorescent protein (YFP) and n = 16 channelrhodopsin-2 (ChR2) male rats. The experiment also included a cohort of female rats with n = 12 each for YFP and ChR2. Throughout the experiments, both male and female rats received the same stimulation and were subjected to real-time place preference (RTPP) and social interaction tests before acute restraint stress and non-invasive measures of hemodynamics. A parallel group of male ChR2 rats (n = 6) was used for slice electrophysiology to verify stimulation parameters. Experiment 2 was comprised of n = 9 green fluorescent protein (GFP) and n = 8 TeNT male rats. A separate female cohort led to n = 9 GFP and n = 9 TeNT. All rats were injected and allowed to recover for 6-8 weeks before assessments. At the end of all experiments, brain tissue was collected to validate the experimental approaches.

**Figure 1:**
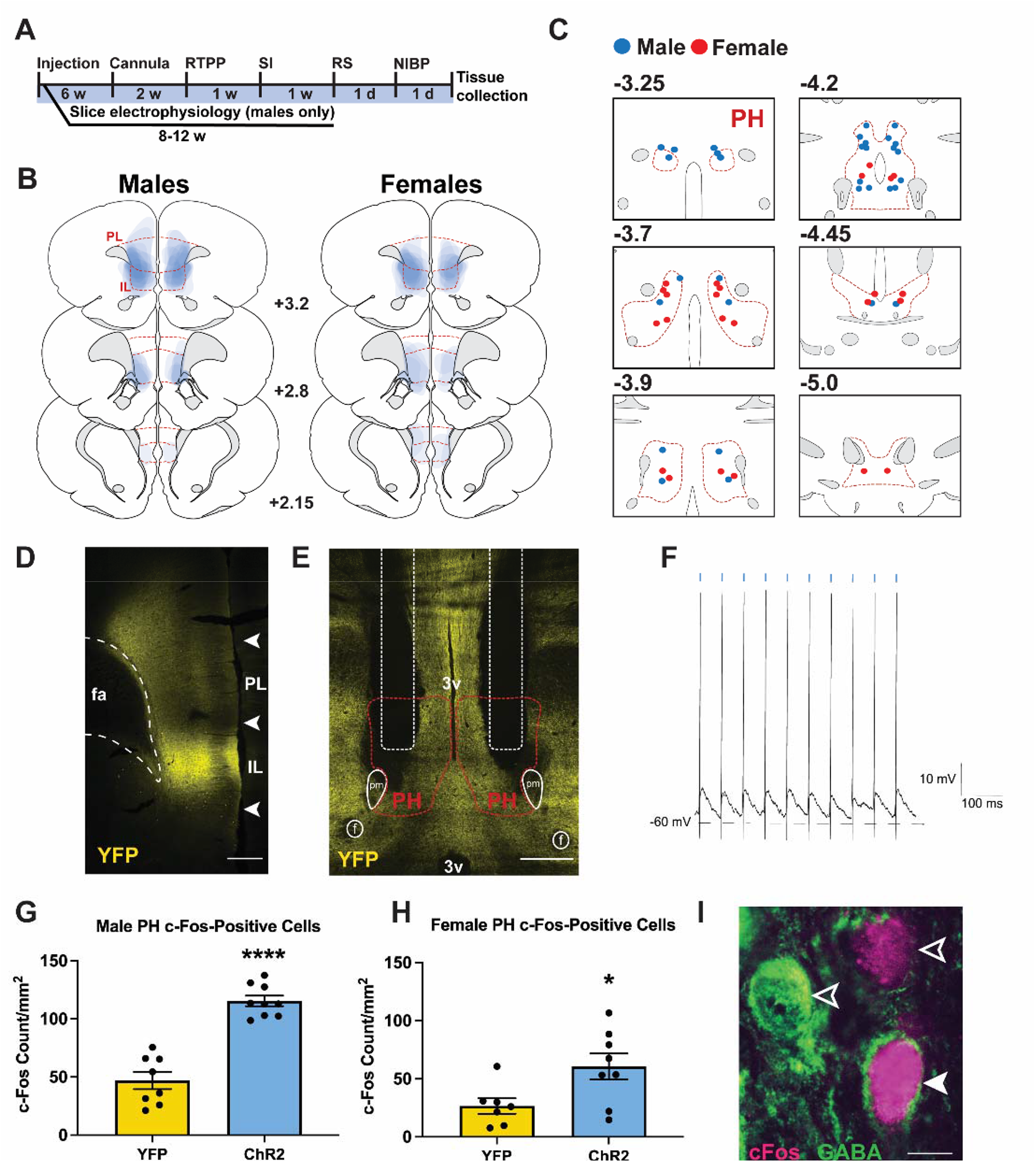
Design and validation for vmPFC-PH stimulation. (**A**) Experimental timeline. w: week, d: day, RTPP: real-time place preference, SI: social interaction, RS: restraint stress, NIBP: non-invasive blood pressure. (**B**) AAV injection spread mapped in blue. Coronal sections adapted from Swanson Rat Brain Atlas (3rd edition). Red lines indicate the border of the infralimbic (IL) and prelimbic (PL) cortices. Numbers indicate distance in mm rostral to bregma. (**C**) Bilateral cannula placements (male: blue, female: red) mapped within the PH (red outline). Numbers indicate distance in mm caudal to bregma. (**D**) Representative photomicrograph of virus spread in the vmPFC (2.8 mm rostral to bregma). Scale bar: 500 μm. (**E**) Representative PH fiber optic cannula placement. White dotted outline: fiber optics, red outline: PH, solid white outline: principal mammillary tract (pm) and fornix (f), 3v: third ventricle. Scale bar: 1000 μm. (**F**) Current-clamp recording of stimulation-locked spiking (10 Hz, 1.1 mW, 5 ms pulse). (**G**) Stimulation increased c-Fos-positive cell density in the male PH (n = 8-9/group) (**H**) as well as the female PH (n = 7-8/group). * p < 0.05, **** p < 0.0001 vs. YFP. (**I**) Representative PH image of c-Fos (magenta) and GABA (green). Unfilled arrows: c-Fos or GABA, white-filled arrows: c-Fos and GABA. Scale bar: 20 μm.

### 2.2 Animals

Age-matched adult male and female Sprague-Dawley rats were obtained from Envigo (Denver, CO) with males ranging from 250-300 g and females 150-200 g. After stereotaxic surgery, rats were individually housed in shoebox cages with cardboard tube enrichment. All rats were housed in temperature- and humidity-controlled rooms with a 12-hour light-dark cycle (lights on at 07:00h, off at 19:00h) and food and water *ad libitum*. In accordance with ARRIVE guidelines, all treatments were randomized, and experimenters were blinded. All procedures and protocols were approved by the Colorado State University Institutional Animal Care and Use Committee (protocol: 1321) and complied with the National Institutes of Health Guidelines for the Care and Use of Laboratory Animals. Signs of poor health and/or weight loss ≥ 20% of pre-surgical weight were a priori exclusion criteria.

### 2.3 Microinjections

For experiment 1, rats were anesthetized with isoflurane (1-5%) followed by analgesic administration (0.6 mg/kg buprenorphine-SR, subcutaneous). Rats received bilateral microinjections (1 µL) of adeno-associated virus (AAV) targeted to the IL portion of vmPFC (Wallace et al., 2021; Wood et al., 2019) (males: 2.7 mm anterior to bregma, 0.6 mm lateral to midline, and 4.0 mm ventral from dura; females: 2.45 mm anterior to bregma, 0.5 mm lateral to midline, and 4.0 mm ventral from dura). AAV5-packaged constructs (University of North Carolina Vector Core, Chapel Hill, NC) expressed either YFP or ChR2 conjugated to YFP under the synapsin promoter to permit terminal stimulation of projection neurons. All microinjections were carried out with a 25-gauge, 2-µL microsyringe (Hamilton, Reno, NV) using a microinjection unit (Kopf, Tujunga, CA) at a rate of 5 minutes/µL. The needle was left in place for 5 minutes before and after injections to reduce tissue damage and allow diffusion. Skin was closed with wound clips that were removed 2 weeks later prior to fiber optic surgery described below. Animals were allowed at least 6 weeks for recovery and ChR2 expression.

For experiment 2, an intersectional approach was used to selectively express tetanus toxin (TeNT) in PH-projecting vmPFC neurons to inhibit neurotransmitter release (Link et al., 1992; McMahon et al., 1993; Sando et al., 2017; Xu and Sudhof, 2013). Using the same surgical procedures described above, rats received bilateral injections (0.75 µL) in the PH of a retrograde AAV-packaged construct expressing Cre. This was followed by vmPFC injections of either 0.75 µL Cre-dependent TeNT or Cre-dependent GFP control. The TeNT virus was obtained from Stanford Gene Vector and Virus Core while the GFP-expressing construct (50457-AAV5) and retrograde-transported Cre (107738-AAVrg) were obtained from Addgene. After injection, rats were allowed 6 weeks for recovery and TeNT expression.

### 2.4 Slice electrophysiology

8 to 12 weeks after injection of ChR2-expressing AAVs in the vmPFC, brain slices were collected for whole-cell recordings as previously described (Rau and Hentges, 2017; Wallace et al., 2021). As detailed in the supplemental methods, optic stimulation parameters were validated for evoking excitatory postsynaptic potentials in the PH.

### 2.5 Fiber optic cannulation

For experiment 1, rats were anesthetized (isoflurane 1-5%) followed by analgesic (0.6 mg/kg buprenorphine-SR, subcutaneous) and antibiotic (5 mg/kg gentamicin, intramuscular) administration before bilateral implantation of fiber optic cannulas (Wallace et al., 2021). Cannulas (flat tip 400/430 μm, NA = 0.66, 1.1 mm pitch with 7.5 mm length; Doric Lenses, Québec, Canada) were targeted to the PH (males and females: 4.0 mm posterior to bregma, 0.5 mm lateral to midline, and 7.5 mm ventral from dura). Cannulas were secured to the skull with metal screws (Plastics One) and dental cement (Stoelting, Wood Dale, IL) before skin was sutured. Following 1 week of recovery, rats were handled daily and acclimated to the stimulation procedure for another week before experiments began. Rat handling and cannula habituation continued daily throughout experiments.

### 2.6 Optogenetic stimulation

For experiment 1, light pulses (8 mW, 5 ms, 10 Hz) were delivered through a fiber-optic patch cord (240 μm core diameter, NA = 0.63; Doric Lenses) connected to a 473 nm LED driver (Doric Lenses). Optic power was measured with a photodiode sensor (PM160, Thorlabs Inc, Newton, NJ) at the cannula fiber tip.

### 2.7 Estrous cycle cytology

Following each assessment, vaginal cytology was examined to approximate the estrous cycle stage. A damp (deionized water) cotton swab was used to collect cells from the vaginal canal and roll them onto a glass slide. Dried slides were viewed under a 10x objective light microscope by a minimum of two blind observers and were categorized as proestrus, estrus, metestrus, or diestrus (Cora et al., 2015; Smith et al., 2018; Solomon et al., 2015).

### 2.8 Real-time place preference

The RTPP assay was used to assess subjects’ preference or aversion for vmPFC-PH stimulation as previously described (Wallace et al., 2021). Methodological details are provided in the supplement.

### 2.9 Social behavior

A modified version of the 3-chambered social behavior assay was used to accommodate optic patch cords as previously reported (Felix-Ortiz and Tye, 2014; Moy et al., 2004; Wallace et al., 2021). Detailed methodologies are outlined in the supplement for testing both social motivation and social novelty preference.

### 2.10 Restraint stress

Restraint was used to examine neuroendocrine responses to acute stress. Blood glucose and plasma corticosterone were measured as indicators of sympathetic (Bialik et al., 1988) and HPA axis (Myers et al., 2014) stress responses, respectively. Rats from both experiments were placed in plastic decapicones (Braintree Scientific, Braintree, MA) with experiment 1 animals connected to fiber optic patch cords for optic stimulation throughout the 30 min restraint as previously described (Wallace and Myers, 2023; Wallace et al., 2021). Once restrained, the distal tail was clipped and blood samples (approximately 250 µL) were collected at the initiation of restraint with additional samples taken 15 and 30 min after (Vahl et al., 2005). After 30 minutes, rats were removed from restraint, patch cords disconnected, and returned to the homecage with additional blood samples collected at 60 and 90 minutes after the initiation of restraint. Blood glucose was determined with Contour Next EZ glucometers (Bayer, Parsippany, NJ) with 2 independent readings for each time point were averaged. Blood samples were centrifuged at 2500 X gravity for 15 min at 4 ^°^C and plasma was stored at −20 ^°^C. Samples were processed in duplicate for corticosterone analysis via Enzo Life Sciences (East Farmingdale, New York) Corticosterone ELISA kit (Catalog No. ADI-900-097) with an intra-assay coefficient of variation of 8.4% and an inter-assay coefficient of variation of 8.2% (Bekhbat et al., 2018; Dearing et al., 2021; Wallace and Myers, 2023).

### 2.11 Non-invasive heart rate and blood pressure recordings

Heart rate (HR) and mean arterial pressure (MAP) were measured to determine whether vmPFC-PH stimulation modulated cardiovascular physiology. Rats were lightly anesthetized with 2% isoflurane then placed prone on a heating pad with anesthesia maintained via nose cone. Body temperature was monitored with a rectal probe and maintained at 37 °C. Cannulas were connected to fiber optic patch cords before a non-invasive blood pressure cuff (Kent Scientific, Torrington, Connecticut) was placed on the base of the tail. The CODA^®^ monitor (Kent Scientific) recorded HR and MAP for 3 minutes before the LED stimulation (473 mm blue light) for 6 minutes. For analysis, data were averaged into 3, 3-min bins (no-stimulation, first 3 mins of stimulation, second 3 mins of stimulation).

### 2.12 Tissue collection

At the conclusion of all experiments, brain tissue was collected for technical validation and immunohistochemical analysis. A subset of experiment 1 rats received 5 min of optogenetic stimulation (8 mW, 5 ms pulses, 10 Hz) 90 minutes prior to tissue collection to examine immediate-early gene (c-Fos) expression. After, rats were injected with sodium pentobarbital (100 mg/kg) and perfused transcardially with 0.9% saline followed by 4.0% paraformaldehyde in 0.1 M phosphate buffer solution (PBS). Brains were removed and post-fixed in 4.0% paraformaldehyde for 24 h at room temperature, followed by storage in 30% sucrose in PBS at 4^°^C. Coronal sections were made on a freezing microtome at 30 μm thickness and then stored in cryoprotectant solution at – 20^°^C.

### 2.13 Immunohistochemistry and microscopy

To verify injection and cannula placement in experiment 1, coronal brain sections were imaged with a Zeiss Axio Imager Z2 microscope using a 10x objective. GABA was immunolabeled to examine potential colocalization of c-Fos and GABA in the PH. For fluorescent labeling of c-Fos and GABA, coronal brain sections were removed from cryoprotectant and rinsed at room temperature in 1X PBS (5 x 5 min). Sections were then submerged into blocking solution (1X PBS, 0.1% bovine serum albumin, and 0.2% Triton X-100) for 1 hour. Then, sections were incubated in polyclonal guinea pig anti-c-Fos primary antibody (Synaptic Systems, Göttingen, Germany; 226-005) (1:1000 in 1X PBS) overnight. The next day, another PBS wash (5 x 5 min) was done before the tissue was incubated in Cy5 donkey anti-guinea pig secondary antibody (Jackson Immuno Research, West Grove Pennsylvania;706-175-148) (1:1000 in 1X PBS) for 1 hour. Following a PBS wash, the tissue was placed into blocking solution (1X PBS, 6% bovine serum albumin, 8% normal goat serum, 2% normal donkey serum, and 0.4% Triton X-100) for 4 hours. The tissue was next incubated in rabbit anti-GABA primary antibody (Sigma, Saint Louis, Missouri; MFCD00162297) (1:250 in blocking solution) for 60 hours at 4 ^°^C. After, the tissue was washed in PBS (5 x 5) and incubated in biotinylated secondary antibody goat anti-rabbit (Vector Laboratories; BA-100) (1:500 in 1X PBS) for 2 hours. Another PBS (5 x 5) wash was conducted before the tissue was incubated in Avidin-Biotin Complex (Vector Laboratories, Newark, California; Vectastain ABC Kit PK-4000) (1:500 in 1X PBS) for 1 hour. The tissue was washed in PBS again (5 x 5) before incubation in Cy3-Streptavidin (Jackson Immuno Research; 016-160-084) (1:500 in PBS) for 1 hour. Finally, the tissue was washed in PBS (5 x 5), mounted, and coverslipped. For quantification of c-Fos -positive cell density in the PH, cells were counted by a treatment-blind observer in 20x-tiled micrographs (10/rat) adjacent to cannula tracts.

For experiment 2, immunohistochemistry was utilized to quantify the percent colocalization of synaptobrevin-2 with GFP in vmPFC projections to the PH. Tissue was removed from cryoprotectant and washed in PBS (5 x 5 mins) before incubation in blocking solution (1X PBS + 0.1% BSA + 0.2% Triton X-100) for 2 hours. After, the tissue was incubated in rabbit anti-GFP primary antibody (Invitrogen, Waltham, Massachusetts; A11122) (1:1000 in blocking solution) overnight. The next morning the tissue was washed in PBS and incubated in A488 goat anti-rabbit secondary antibody (Invitrogen, A11008) (1:500 in 1X PBS) for 30 mins. Following a PBS wash, tissue was blocked again (4% BSA, 3% donkey serum, 0.1% Triton) for 1 hour before incubation in synaptobrevin-2 primary antibody (Synaptic Systems; 104 211C3) (1:200 in blocking solution) overnight at 4^°^C. On the final day, the tissue was washed in PBS (5 x 5) and incubated in Cy5 donkey anti-mouse secondary (Jackson Immuno Research; 715-175-150) (1:500 in PBS) for 2 hours. Tissue was then washed in PBS (5 x 5) and mounted for microscopy. Slides were imaged with a Zeiss Axio Imager Z2 microscope. On average, 6 images of PH-projecting vmPFC axons were taken per rat. 5-image z-stacks, spanning 2.5 μm, were captured using a 63X objective with apotome processing. Using Zeiss colocalization tool (Zeiss Blue edition version Zen 2.6 pro), a blinded observer applied a colocalization threshold to individual axons in the PH to calculate the percentage of the GFP axon colocalized with Cy5 fluorescence labeling synaptobrevin-2 protein.

### 2.14 Data analysis

All data are presented as mean ± standard error of the mean. Data were analyzed using Prism 9.3.1 (GraphPad, San Diego, CA), with statistical significance set at *p* < 0.05 for rejection of null hypotheses. Stimulation-induced c-Fos and RTPP side preference were analyzed with Welch’s unpaired t-tests comparing virus groups (YFP vs. ChR2). Social motivation and novelty preference were assessed with unpaired t-tests to compare virus groups (YFP vs. ChR2) within social, object, novel rat, or familiar rat interactions. Total distance traveled during behavioral testing was also assessed with unpaired t-tests comparing virus groups. Time-dependent responses (corticosterone, glucose, and non-invasive hemodynamics) were analyzed using mixed-effects analysis with virus (YFP vs. ChR2 or GFP vs. TeNT) and time (repeated) as factors, followed by Fisher’s post-hoc if significant main or interaction effects were present. Synaptobrevin-2 colocalization with GFP was analyzed with unpaired t-tests comparing virus groups (GFP vs. TeNT).

## 3. Results

### 3.1 Optogenetic stimulation of vmPFC-PH circuit

AAV viral vectors were injected into the vmPFC to express YFP or ChR2-YFP. Mapped viral spread (**Fig. 1B**) shows injections targeted to the IL with some dorsal spread to the prelimbic cortex (PL) with similar coverage in both sexes. Mapped fiber optic cannula placements (**Fig. 1C**) show fibers were targeted to the PH. Representative photomicrographs depict virus injection in the vmPFC and bilateral cannula placement in the PH (**Fig. 1D&E**). To validate stimulation parameters, whole-cell patch-clamp recordings from PH slices indicated that 10 Hz optogenetic stimulation produced excitatory postsynaptic potentials (**Fig. 1F**). *In vivo* stimulation at 10 Hz also increased PH expression of the immediate-early gene c-Fos approximately 2.5-fold in both ChR2 males (**Fig. 1G**; males: n = 8-9/group, unpaired t-test: t(11.89) = 7.90, *p* < 0.0001) and females (**Fig. 1H**; females: n = 7-8/group, unpaired t-test: t(11.21) = 2.62, *p* = 0.024) compared to YFP controls. Additionally, stimulation-activated PH cells in males and females were both GABAergic and non-GABAergic (**Fig. 1I**).

### 3.2 Real-time place preference

RTPP was used to examine the affective valence of stimulating vmPFC terminals in the PH. Group mean heat maps illustrate time spent in the stimulation versus non-stimulation chambers (**Fig. 2A**). Male ChR2 rats preferred the stimulation chamber compared to YFP (**Fig. 2B**; males: n = 15-16/group, unpaired t-test: t(21.34) = 5.50, *p* < 0.0001). There was no stimulation preference or aversion in female rats. (**Fig. 2C**; females: n = 12/group, unpaired t-test: t(13.37) = 1.42, *p* = 0.18). Additionally, there were no stimulation effects on overall locomotor behavior in either sex. Results indicate that activity of the vmPFC-PH circuit induces positive valence in male, but not female, rats.

**Figure 2:**
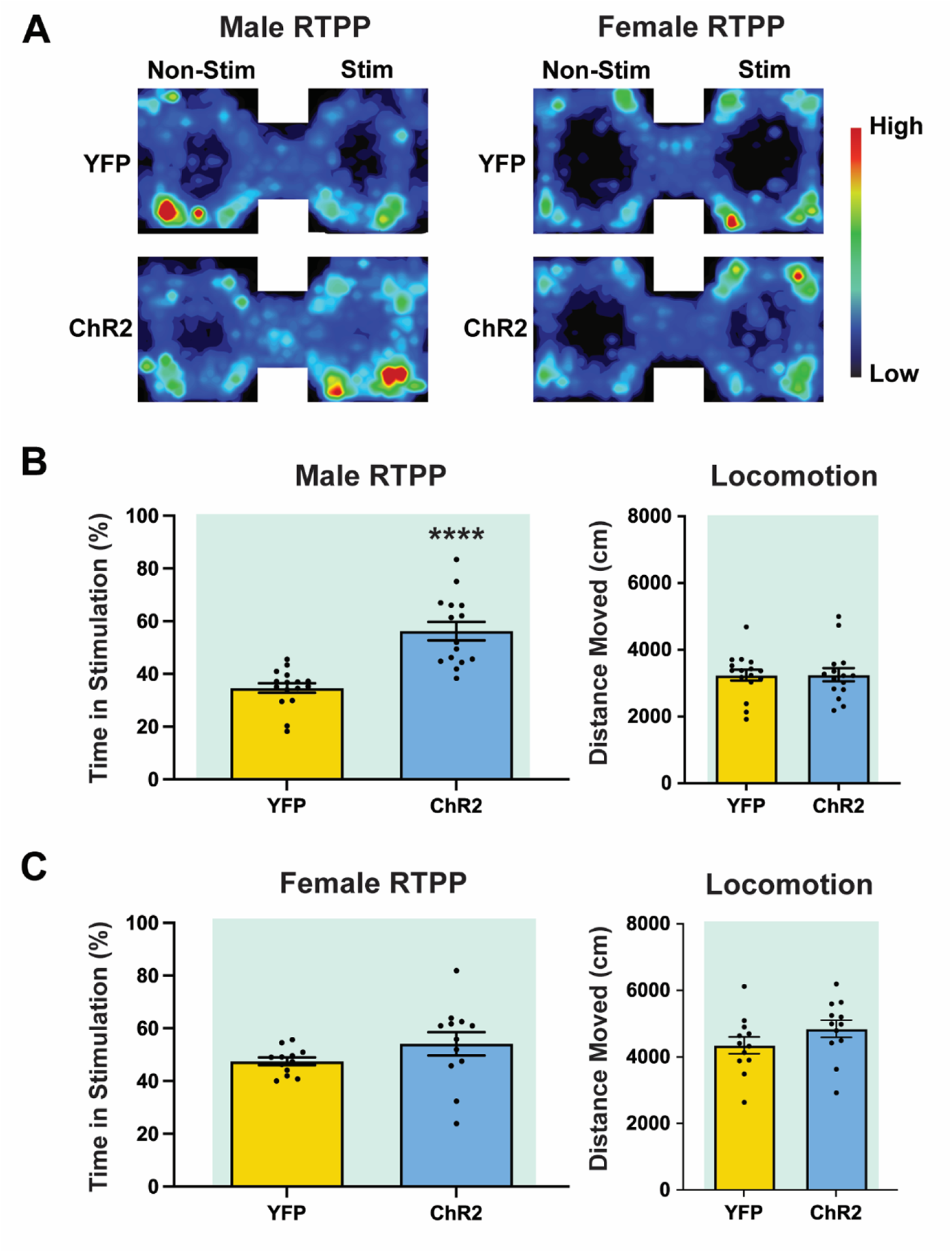
Stimulation of the vmPFC-PH circuit had positive valence in males only. (**A**) Heat maps display mean animal position in the RTPP arena. (**B**) Male ChR2 rats preferred the chamber paired with stimulation relative to YFP controls. Total distance traveled found no treatment-based differences in locomotion. (**C**) Female ChR2 rats had no preference or aversion for the stimulation chamber relative to YFP controls and no difference in locomotion. n = 10-15/group, **** p < 0.001 vs. YFP.

### 3.3 Social behavior

The three-chamber social interaction assay was used to determine the influence of vmPFC-PH stimulation on sociability. There were no differences in social motivation in either sex (**Table S1**; males: n = 15/group, unpaired t-test: t(24.55) = 0.10, *p* = 0.92; females: n = 12/group, unpaired t-test: t(19.09) = 0.35, *p* = 0.73). Further, stimulation did not alter social novelty preference or total distance traveled in either sex. These results indicate that vmPFC inputs to the PH are not sufficient to alter social behavior.

### 3.4 Endocrine reactivity: vmPFC-PH stimulation

To determine the sufficiency of vmPFC-PH signaling for sympathetic and HPA axis responses, blood glucose and plasma corticosterone were analyzed during restraint stress. In males, optic stimulation decreased glucose mobilization (**Fig. 3A**; n = 16-17/group, mixed-effects: time F(4,119) = 34.59, *p* < 0.0001, ChR2 F(1,31) = 1.35, p = 0.25, time x ChR2 F(4,119) = 1.16, *p* = 0.33) during restraint (15 min, *p* = 0.021), but did not affect blood glucose in females (**Fig. 3B**; n = 12/group, mixed-effects: time F(4,78) = 24.63, *p* < 0.00001, ChR2 F(1,21) = 0.18, *p* = 0.67, time x ChR2 F(4,78) = 0.45, *p* = 0.77). Stimulation did not affect male plasma corticosterone (**Fig. 3C**; n = 16-17/group, mixed effects: time F(4,114) = 74.15, *p* < 0.0001, ChR2 F(1,31) = 0.48, *p* = 0.50, time x ChR2 F(4,114) = 0.42, *p* = 0.79), but stimulation increased plasma corticosterone in female rats (**Fig. 3D**; n = 12/group, mixed effects: time F(4,99) = 23.91, *p* < 0.0001, ChR2 F(1,99) = 3.49, *p* = 0.065, time x ChR2 F(4,99) = 2.32, *p* = 0.062) at the 60- and 90-minute timepoints (60 min, *p* = 0.0049, 90 min, *p* = 0.035). Collectively, these results suggest that vmPFC projections to the PH reduce male sympathetic activity but are not sufficient to alter corticosterone secretion. In females, circuit stimulation does not alter glucose mobilization but increases HPA axis activity.

**Figure 3:**
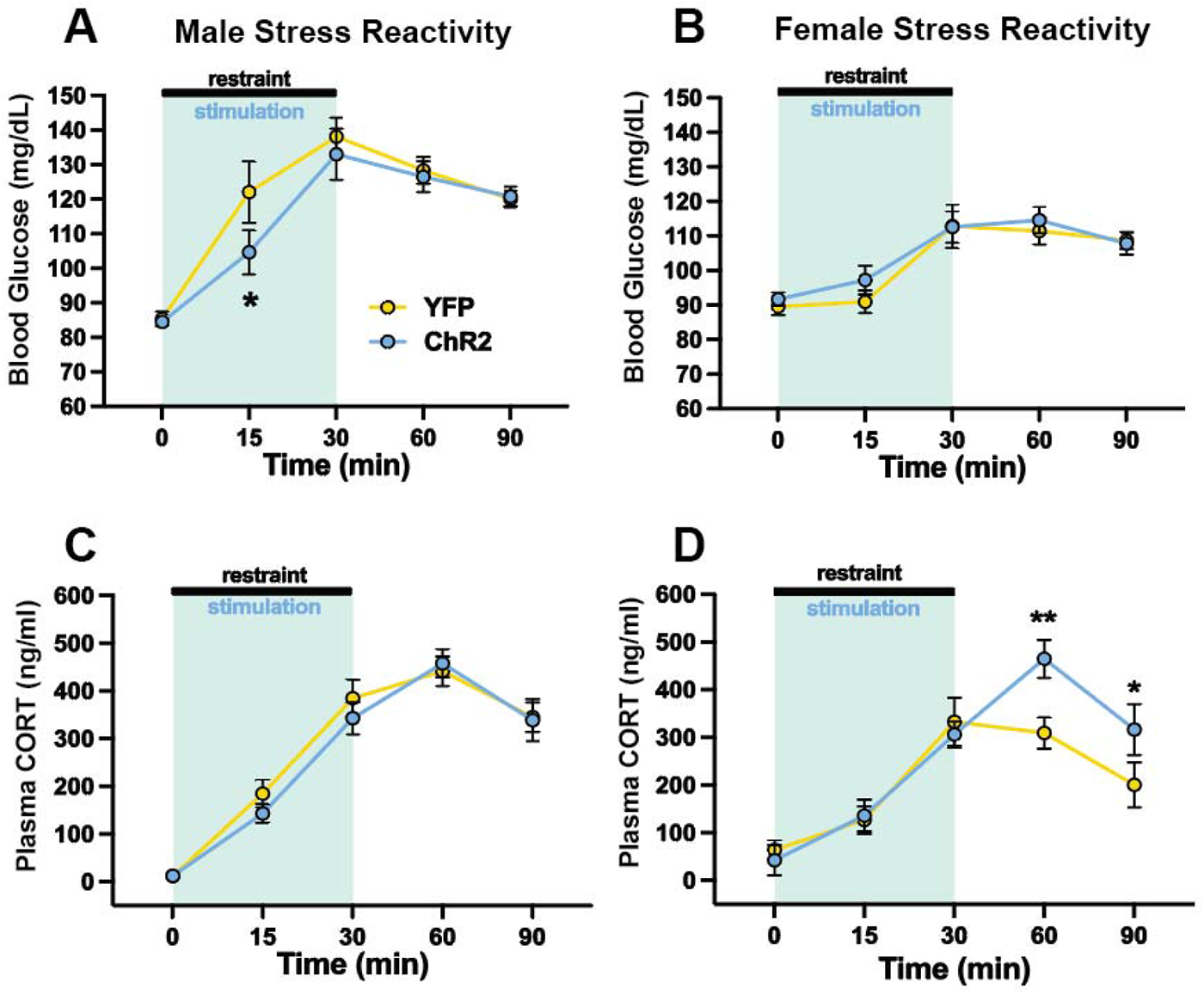
Stimulation of vmPFC-PH projections reduced male glucose mobilization but increased female corticosterone responses to restraint stress. (**A**) Blue light stimulation (blue shading) decreased blood glucose in ChR2 males compared to YFP controls during restraint. (**B**) Stimulation had no effect on blood glucose in females. (**C**) Stimulation did not affect plasma corticosterone (CORT) in males but increased CORT responses in females (**D**) following restraint. n = 10-17/group, * p < 0.05, ** p < 0.01 vs. YFP.

### 3.5 Hemodynamics

To determine whether vmPFC-PH stimulation influences cardiovascular parameters, a non-invasive tail cuff was used to measure HR and MAP during optic stimulation. In males, acute stimulation had no effect on HR (**Fig. 4A**; n = 5-6/group, mixed-effects: time F(2,16) = 0.24, *p* = 0.79, ChR2 F(1,9) = 0.66, *p* = 0.44, time x ChR2 F(2,16) = 0.0055, *p* = 0.995). In females, stimulation increased HR (**Fig. 4B**; n = 7-10/group, mixed effects: time F(2,20) = 0.67, *p* = 0.53, ChR2 F(1,15) = 6.64, *p* = 0.021, time x ChR2 F(2,20) = 3.75, *p* = 0.041) during the first 3 minutes of stimulation (3 min, *p* = 0.009, 6 min, *p* = 0.053). Stimulation had no effects on male (**Fig. 4C**; n = 7-8/group, mixed effects: time F(2,18) = 13.56, *p* < 0.001, ChR2 F(1,13) = 3.16, *p* = 0.099, time x ChR2 F(2,18) = 0.82, *p* = 0.45) or female MAP (**Fig. 4D**; n = 9-12/group, mixed effects: time F(2,25) = 8.99, *p* < 0.01, ChR2 F(1,19) = 0.095, *p* = 0.76, time x ChR2 F(2,25) = 0.11, *p* = 0.90). Collectively, anesthetized hemodynamic recordings indicate that vmPFC-PH stimulation elevates HR specifically in females.

**Figure 4:**
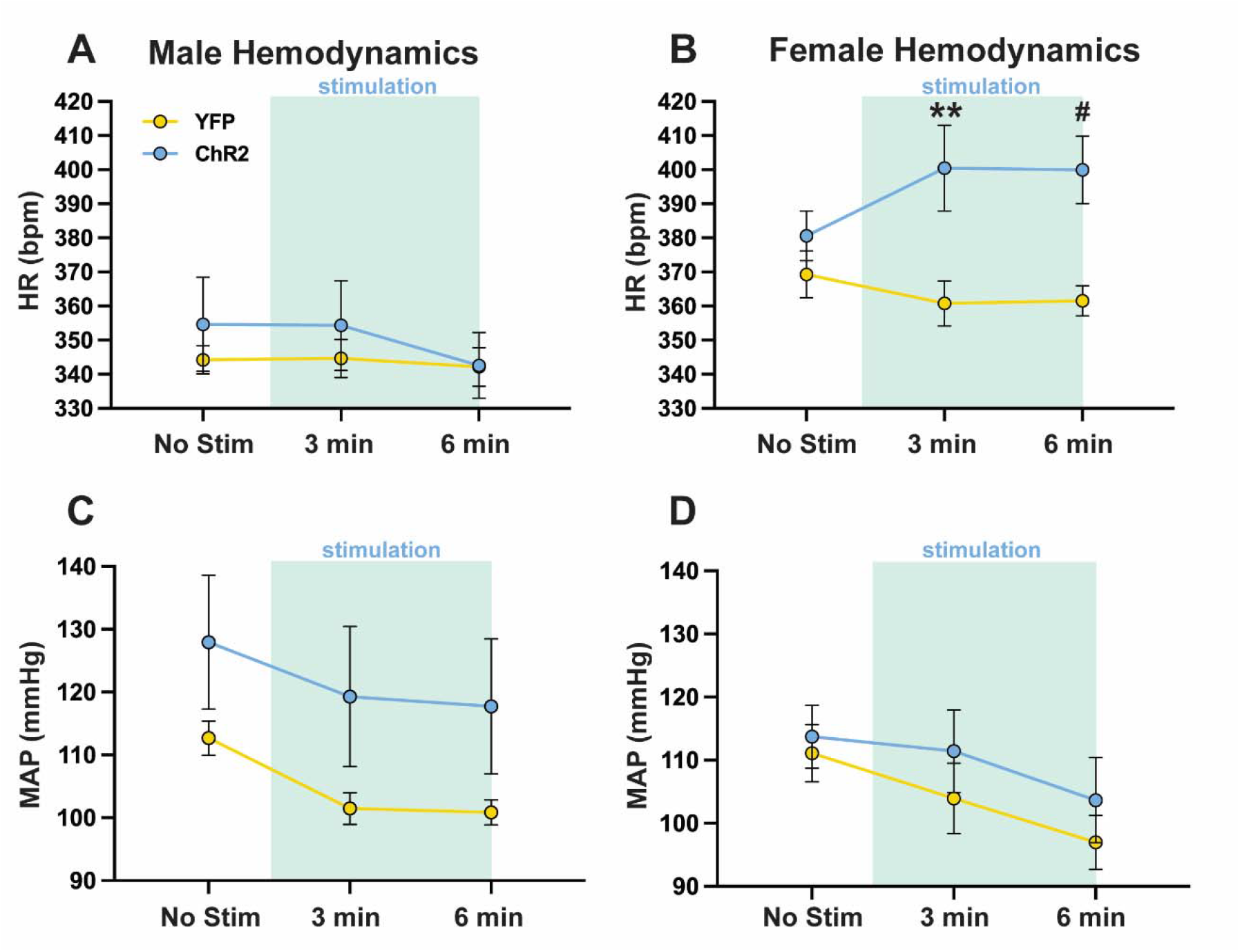
vmPFC-PH stimulation elevated heart rate in females. (**A**) Stimulation (blue shading) during non-invasive tail cuff recordings had no effect on heart rate (HR) in males. (**B**) Stimulation elevated HR in females compared to YFP controls. Stimulation had no effect on blood pressure in males (**C**) or females (**D**). bpm: beats per minute. All rats were maintained under 2% isoflurane during hemodynamic recordings. n = 5-8/group, ** *p* < 0.01, ^#^ *p* = 0.053, vs. YFP.

### 3.6 Inhibition of PH-projecting vmPFC neurons

Intersectional genetics was used to test the necessity of the vmPFC-PH circuit for sympathetic and HPA axis stress responses. PH-targeted microinjections of a retrograde-transported AAV carrying a construct for Cre expression were mapped on coronal sections adapted from (Swanson, 2004) (**Fig. 5A**). Representative micrographs of dTomato-labelled retrograde-Cre injected in the PH (**Fig, 5B**) and Cre-dependent GFP expression in the vmPFC (**Fig. 5C**) indicate successful recombination following vmPFC and PH injections. Representative micrographs illustrate synaptobrevin-2 expression on GFP-positive axons in the PH of both GFP-(**Fig. 5D**) and TeNT-injected (**Fig. 5E**) rats. Colocalization quantification found that TeNT significantly reduced (by approximately 70%) the percentage of GFP axonal length expressing synaptobrevin-2 protein in both males (**Fig. 5F**; n = 6-7/group, unpaired t-test: t(10.46) = 9.40, *p* < 0.0001) and females (**Fig. 5G**; n = 7-8/group, unpaired t-test: t(7.48) = 9.64, *p* < 0.0001). As synaptobrevin-2 is essential for neurotransmitter exocytosis, this approach has been used to reduce synaptic communication (Link et al., 1992; Sando et al., 2017; Xu and Sudhof, 2013).

**Figure 5:**
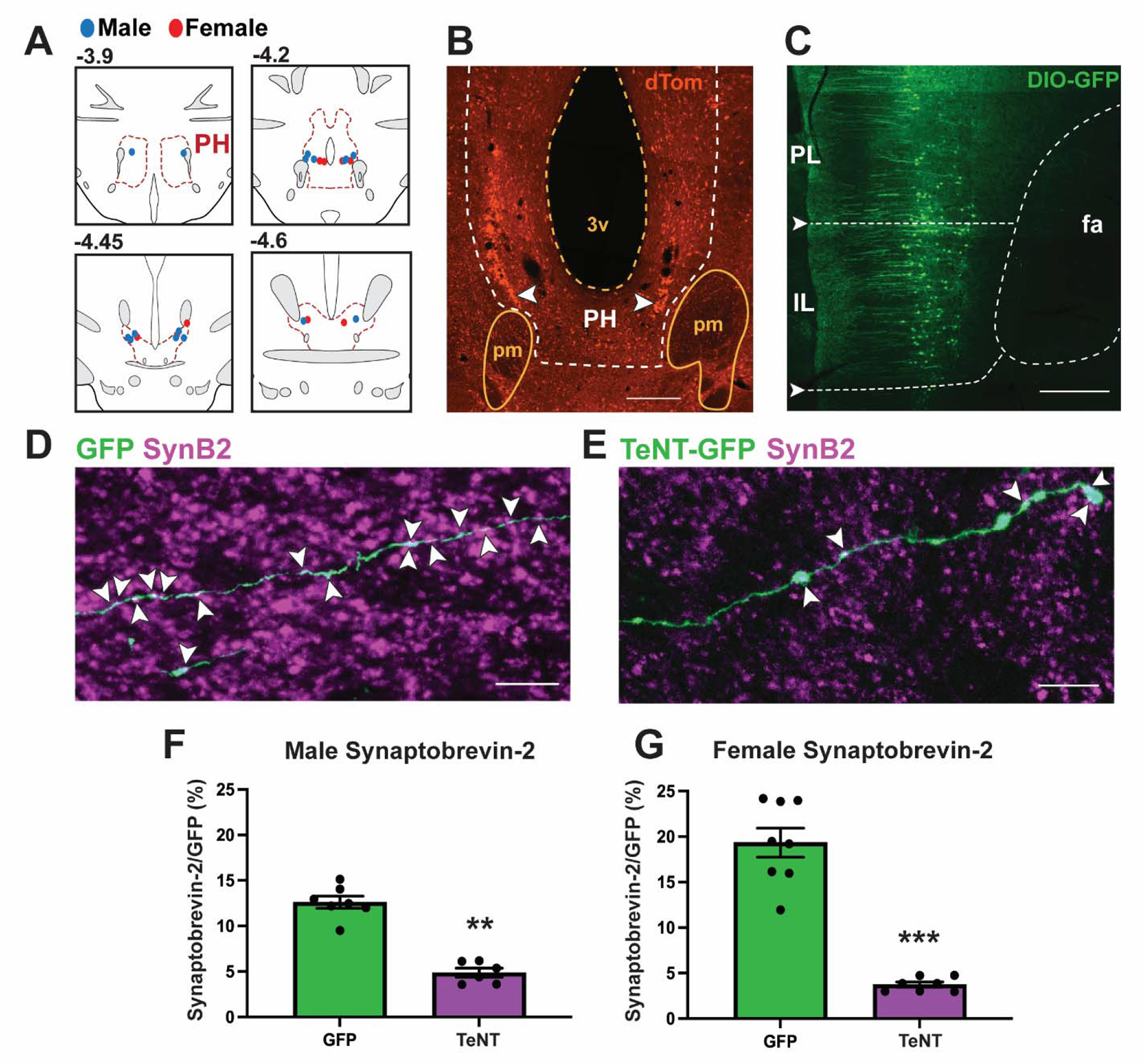
Synaptobrevin knockdown in PH-projecting vmPFC neurons. (**A**) Bilateral injections of retrograde AAV carrying Cre construct. Coronal sections adapted from the Swanson Rat Brain Atlas (Swanson, 2004). Numbers indicate the distance in mm caudal to bregma. Male and female injections are labeled with blue and red dots, respectively. (**B**) Representative photomicrograph of retrograde-Cre virus injected into the PH. Scale bar: 200 μm. (**C**) Representative photomicrograph demonstrates expression of Cre-dependent GFP targeted to the IL with limited spread to the PL. fa: anterior forceps of the corpus callosum. Scale bar: 200 μm. Representative micrographs of synaptobrevin-2 protein (SynB2; magenta) colocalizing (white) with GFP on vmPFC axons in the PH of GFP-(**D**) and TeNT-treated (**E**) rats. Scale bar: 20 μm. Synaptobrevin-2 protein was decreased in both male (**F**) and female (**G**) TeNT-treated rats compared to GFP controls. n = 6-8/group, ** *p* < 0.01, *** *p* < 0.001, vs GFP.

### 3.7 Endocrine reactivity: vmPFC-PH inhibition

To test the necessity of PH-projecting vmPFC neurons for sympathetic and HPA axis stress responses, blood samples were collected to measure blood glucose and plasma corticosterone. vmPFC-PH circuit inhibition elevated blood glucose mobilization in males (**Fig. 6A**; n = 9/group, mixed effects: time F(4,64) = 20.9, *p* < 0.0001, TeNT F(1,16) = 2.79, *p* = 0.11, time x TeNT F(4,64) = 3.49, *p* = 0.012; 15 min, *p* = 0.023; 30 min, *p* = 0.002) but not females (**Fig. 6B**; n = 9/group, mixed effects: time F(4,68) = 2.05, *p* = 0.097, TeNT F(1,18) = 1.63, *p* = 0.22, time x TeNT F(4,68) = 0.92, *p* = 0.46).

**Figure 6:**
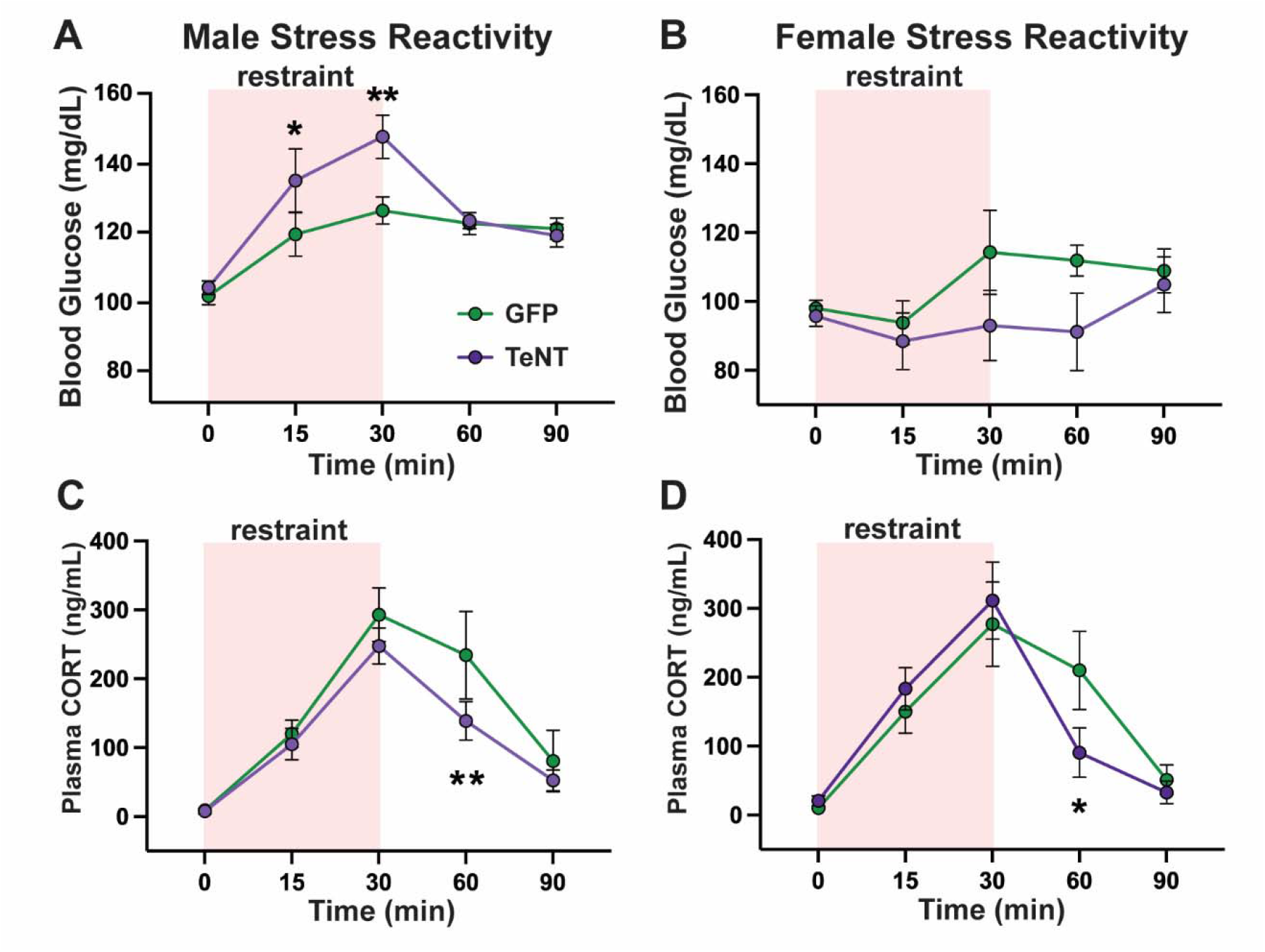
Inhibition of PH-targeting vmPFC neurons increased glucose mobilization in males and decreased corticosterone responses to restraint stress in both sexes. IL-PH circuit inhibition elevated blood glucose mobilization in males **(A)** but not females **(B)**. Circuit inhibition enhanced recovery of corticosterone responses in males **(C)** and females **(D)**. n = 8-9/group, * p < 0.05, ** p < 0.01 vs. GFP.

TeNT reduced plasma corticosterone on male (**Fig. 6C**; n = 8/group, mixed effects: time F(4,23) = 13.40, *p* < 0.0001, TeNT F(1,14) = 3.52, *p* = 0.082, time x TeNT F(4,23) = 1.60, *p* = 0.21; 60 min, *p* = 0.0038) and female HPA axis reactivity (**Fig. 6D**; n = 9/group, mixed effects: time F(4,49) = 20.10, *p* < 0.0001, TeNT F(1,17) = 0.22, *p* = 0.64, time x TeNT F(4,49) = 1.46, *p* = 0.23) at the 60 minute timepoint (60 min, *p* = 0.042).

Altogether, TeNT-mediated circuit inhibition in males enhanced blood glucose during acute restraint yet attenuated plasma corticosterone in both sexes.

### 3.8 Estrous cycle cytology

The distribution of rats in each estrous phase is included in **Tables S2** and **S3**. Within-phase sample sizes did not permit further statistical analysis.

## 4. Discussion

In the current study, circuit-specific approaches were used in male and female rats to examine the role of vmPFC-PH projections in behavior, stress reactivity, and cardiovascular regulation. Stimulation of vmPFC-PH terminals increased preference behavior and inhibited stress-induced glucose mobilization in males. Furthermore, inhibition of PH-targeting vmPFC neurons in males during acute stress increased glucose mobilization but also decreased corticosterone responses. In females, stimulation of the vmPFC-PH circuit did not affect behavior or glucose homeostasis but increased corticosterone stress responses and elevated HR. Additionally, inhibition of the female vmPFC-PH circuit reduced corticosterone stress responses. Collectively, vmPFC-PH circuitry functions in a sexually-divergent manner where male PH-projecting vmPFC neurons are sufficient to influence behavior and reduce sympathetic-mediated stress responses, while the female circuit is sufficient to increase corticosterone secretion and HR. Additionally, the male and female vmPFC-PH circuit is necessary for increasing glucocorticoid stress responses, yet the male circuit is necessary to inhibit glucose stress responses.

In rodent models, the male vmPFC modulates HPA axis (McKlveen et al., 2013; Radley et al., 2006) and autonomic cardiovascular responses to stress (Müller-Ribeiro et al., 2012; Resstel et al., 2004). Furthermore, male vmPFC pyramidal glutamate neurons regulate acute cardiovascular stress reactivity, as well as the cardiac, vascular, and behavioral consequences of chronic stress (Pace et al., 2020; Schaeuble et al., 2019; Wallace et al., 2021). Previously, we found that stimulation of vmPFC glutamate neurons increases place preference and social motivation in male but not female rats (Wallace et al., 2021). Data from the current study indicate that male preference behaviors are mediated by vmPFC projections to the PH. However, vmPFC synapses in the PH did not modulate social behavior, suggesting the vmPFC may regulate social behavior through synapses in other regions such as the amygdala (Felix-Ortiz and Tye, 2014). Furthermore, vmPFC pyramidal neuron stimulation in males broadly inhibits physiological responses to stress (Wallace et al., 2021). Here, we found that vmPFC-PH circuitry partially accounted for male cortical stress inhibition as the pathway was sufficient and necessary to inhibit stress-induced glucose mobilization. However, vmPFC-PH stimulation did not affect male corticosterone stress responses and circuit inhibition facilitated HPA axis stress recovery. In contrast to males, female vmPFC stimulation enhances stress responding (Wallace et al., 2021). The current data indicate that female vmPFC-PH circuitry increases anesthetized HR and is both sufficient and necessary to increase corticosterone stress responses. Altogether, the data suggest that the PH is a pivotal circuit node that can partially account for sex-specific cortical stress regulation.

Importantly, vmPFC projection neurons synapse onto GABAergic neurons in the PH and GABA_A_ receptor-mediated signaling in the male PH inhibits physiological and behavioral threat responses (Myers et al., 2016; Shekhar et al., 1990). Moreover, the PH projects separately to neuroendocrine and sympathetic nuclei (Nyhuis et al., 2016), suggesting these processes may be distinctly regulated. Little has been reported about the female PH, but the current study indicates cortical inputs to the PH produce sex-specific responses to stress. Although the mechanisms that account for sex-biased circuit function remain to be determined, several possibilities exist. Projections from the vmPFC to the PH are primarily glutamatergic (Myers et al., 2014), yet the sex-specific postsynaptic signaling in the PH is unknown. Male PH glutamate output is gated by local GABA-mediated inhibition (Abrahamson and Moore, 2001; DiMicco and Abshire, 1987; Myers et al., 2016; Nyhuis et al., 2016; Ulrich-Lai et al., 2011; Wible et al., 1988). Here, we found that male and female vmPFC-PH stimulation led to PH c-Fos expression in both GABAergic and non-GABAergic cells. Thus, biological sex may influence the proportion or magnitude of vmPFC synaptic inputs to GABAergic and non-GABAergic cells. PH outputs may also be sexually differentiated. The PH is interconnected with surrounding hypothalamic nuclei, cortical and limbic systems, and brainstem homeostatic regions (Abrahamson and Moore, 2001; Schaeuble and Myers, 2022; Ulrich-Lai et al., 2011; Vertes and Crane, 1996; Ziegler et al., 2002). However, reports of PH circuitry come exclusively from male rodents; therefore, circuit function may be influenced by sex differences in the output of vmPFC-innervated PH cells. Additionally, sex hormones may modulate vmPFC-PH circuit signaling. Estrogen receptors α, β (ERα and ERβ), and G-protein coupled estrogen receptor (GPER) are present in the vmPFC (Almey et al., 2014; Montague et al., 2008; Wallace et al., 2021) where estradiol modulates the excitability of vmPFC neurons and influences behavior (Almey et al., 2014; Dossat et al., 2018; Yousuf et al., 2019). ER mRNA and protein are also expressed throughout the hypothalamus and ER signaling modulates HPA axis activity (Dearing et al., 2022; Handa and Weiser, 2014; Kudwa et al., 2014; Laflamme et al., 1998; Simerly et al., 1990). Additionally, androgen receptors are expressed in cortex and hypothalamus (Aubele and Kritzer, 2012; DonCarlos et al., 2006) and may contribute to differences in stress responding (Handa and Weiser, 2014). Aromatase is also expressed throughout the cortex and hypothalamus, potentially affecting levels of estrogens and androgens (Lu et al., 2019; Wei et al., 2014). Moreover, androgens and estrogens have contrasting effects on vmPFC neurotransmitter metabolism during acute stress (Handa et al., 1997). Altogether, sex-specific vmPFC-PH circuit effects may be due to differences in synaptic excitatory/inhibitory signaling, circuit connectivity, gonadal hormones, and/or brain-derived hormones.

While these experiments identify a novel cortical-hypothalamic pathway for sex-specific stress integration, there are some interruptive limitations. For this study, AAV injections were targeted to the IL, the ventral subregion of the vmPFC. Due to sparse but consistent dorsal injection spread into the PL, we termed the targeted region vmPFC for accuracy. Although there are reports that IL and PL lesions have opposing effects on stress reactivity (Radley et al., 2006; Tavares et al., 2009), IL innervation of the PH is more robust and stress-responsive (Abrahamson and Moore, 2001; Myers et al., 2016; Vertes, 2004). The experiment 2 approach reduced synaptobrevin-2 in both males and females suggesting that the protein was cleaved, thus inhibiting vesicle fusion to the presynaptic membrane. However, the extent of neurotransmission inhibition was not directly accessed. Yet, numerous reports indicate that similar approaches to reduce synaptobrevin lead to a loss of evoked responses (Boehringer et al., 2017; Han et al., 2015; Sando et al., 2017). Another limitation involves the use of anesthesia during hemodynamic recordings. This was necessary to achieve the non-invasive recordings in tethered rats. However, the absence of effects under sedation does not rule out a role for vmPFC-PH pathway regulation of cardiovascular stress responses. In fact, our prior work with vmPFC stimulation found that cardiovascular stress reactivity was modulated in the absence of effects on baseline HR or MAP (Wallace et al., 2021). The specificity of experimental approaches is another important consideration. Experiment 1 used an optogenetic approach with light delivery in the PH for circuit-specific examination. However, experiment 2 used an intersectional genetic approach to knockdown synaptobrevin-2 in PH-projecting vmPFC neurons. The retrograde nature of this method could lead to inhibition of neurotransmitter release from PH-projecting vmPFC neurons at other synaptic targets. Although these approaches have differences in specificity and timing of circuit regulation, they largely generated complementary results.

In conclusion, the current data revealed that stimulating vmPFC synapses in the PH yields sexually divergent outcomes on behavior and stress responding. Moreover, reduced activity of vmPFC projections to the PH also support a role for the circuit in sex-specific stress regulation. Taken together, vmPFC-PH signaling may explain sex-based differences in stress susceptibility.

## Funding

This work was supported by National Institutes of Health grant R01 HL150559 to Brent Myers and American Heart Association Predoctoral Fellowship 827519 to Derek Schaeuble.

## Acknowledgments

AAV5-hSyn-YFP and AAV5-hSyn-ChR2 vectors were provided by the University of North Carolina Vector Core under a material transfer agreement with Karl Deisseroth and Stanford University. AAV5-hSyn-EGFP was a gift from Bryan Roth (Addgene prep # 50457-AAV5) and AAVrg-hSyn-Cre-P2A-dTomato was a gift from Rylan Larsen (Addgene prep # 107738-AAVrg). Additionally, the authors wish to thank Carlie McCartney and Carley Dearing for assistance with sample collection and Dr. Courtney Bouchet and Dr. Adam Chicco for helpful comments on the manuscript.

## Disclosures

The authors have no conflicts of interest to disclose.

## Supplemental Methods

### Slice electrophysiology

Brain sections were collected in ice-cold artificial CSF (aCSF) consisting of the following (in mM): 126 NaCl, 2.5 KCl, 1.2 MgCl_2_ _ 6H_2_O, 2.4 CaCl_2_ _ 2H_2_O, 1.2 NaH_2_PO_4_, 11.1 glucose, and 21.4 NaHCO_3_, bubbled with 95% O_2_ and 5% CO_2_. Coronal slices containing the PH were cut at a thickness of 240 μm using a model VT1200S vibratome (Leica Microsystems, Buffalo Grove, IL). After resting for 1 hour at 37 °C in aCSF, slices were transferred to the recording chamber and perfused with oxygenated 37 °C aCSF at a 2 ml/min flow rate. For whole-cell recordings, the internal recording solution contained the following (in mM): KCl 57.5, K-methyl sulfate 57.5, NaCl 20, MgCl_2_ 1.5, HEPES 5, EGTA 0.1, ATP 2, GTP 0.5, and phosphocreatine 10. The pH was adjusted to 7.3. Recording electrodes had a resistance of 2 – 4 MΩ when filled with this solution. Neurons were identified for recording based on the expression of anterograde ChR2-YFP near the cell. Whole-cell patch-clamp recordings were acquired in current-clamp while holding the current at 0 pA using an Axopatch 200B Amplifier (Molecular Devices, San Jose, CA). Electrophysiological data were collected and analyzed using Axograph X software on a Mac OS X operating system (Apple, Cupertino, CA). Light activation of vmPFC synapses expressing ChR2 was triggered via a 473 nm LED (1.1 mW, Thorlabs, Newton, NJ) light pulse driven by a LEDD1B driver (Thorlabs, Newton, NJ) triggered through the TTL output on an ITC-18 computer interface board (HEKA Instruments, Holliston, MA). Voltage-clamp experiments used a 100 ms light pulse while current-clamp experiments used 5 ms pulses. Current-clamp experiments utilized 10 and 20 Hz stimulation frequencies for 5 min bouts. 10 Hz stimulation generated greater spike fidelity than 20 Hz and was used for all subsequent experiments. Recordings were excluded if access resistance exceeded 10 Ω during recording.

### Real-time place preference

Rats were briefly handled while LED patch cords were connected to cannulas for light delivery. Next, rats were placed in a custom-made fiberglass arena with two chambers connected by a corridor for 10 minutes on two consecutive days (chambers: 15 × 15’’, corridor: 8 × 6’’, 15’’ deep). On the first day, no stimulation was delivered on either side. On the second day, rats received 470 nm light throughout the time spent in the stimulation side. Stimulation stopped when rats exited the assigned stimulation side but re-commenced upon re-entry. Thus, rats determined the amount of stimulation received through time spent in the stimulation side. Trials were recorded by a camera mounted above the arena and animal position was tracked by Ethovision software (Noldus Information Technologies) for automated optic hardware control. The stimulation side assignment was counter-balanced and animal testing was randomized. The time rats spent in the stimulation side was divided by the total time and multiplied by 100 to generate a percentage of time spent in the stimulation side. Total distance traveled was also recorded as a measure of locomotion.

### Social behavior

Each rat was connected to a patch cord and placed in a black rectangular fiberglass arena (36 x 23’’, 15.8’’ deep). Initially, the arena was empty and experimental rats were allowed to explore for 5 minutes without optic stimulation. Next, the experimental rat was returned to the homecage while an empty enclosure (ventilated with small round openings) was placed on one side of the arena, defined as the object, and an identical enclosure containing an age- and sex-matched conspecific was placed on the other side of the arena, defined as the social cage. Again, the experimental rat was then placed in the middle of the arena and allowed to explore freely for 10 minutes with pulsatile optic stimulation delivered throughout to quantify time interacting with the object or social cage as a measure of social motivation. Last, the experimental rat was placed again into the homecage while the empty enclosure was replaced with a new enclosure containing a novel age- and sex-matched conspecific, defined as the novel social cage. The experimental rat freely explored for 10 minutes while receiving optic stimulation to assess social novelty preference. Behavior was recorded with an overhead camera and interactions were defined as nose pokes onto cages and scored by a treatment-blinded observer. The duration of interactions was divided by the total time of each interaction period and multiplied by 100 to give a percent interaction value. Sides for the object cage, social cage, and novel social cage were counterbalanced and animal order randomized.

## Supplemental Data

**Table 1:** Social Behavior. **Stimulation of the vmPFC-PH circuit did not affect social motivation or social novelty preference in males or females.** Time interacting, defined as a nose-poke onto the cage, was not altered by ChR2 stimulation in the object vs. social or novel social vs. familiar social paradigms. Total locomotion was also not altered by ChR2. Values are expressed as mean ± standard error of the mean. n = 12-16/group

|  |  |  |  | Novel | Familiar | Object/Social | Novel/Familiar |
| --- | --- | --- | --- | --- | --- | --- | --- |
|  |  | Object (s) | Social (s) | Social (s) | Social (s) | Distance (cm) | Distance (cm) |
| **Males** | **YFP** | 80.3 ± 11.3 | 320.9 ± 17.8 | 197.3 ± 21.6 | 160.1 ± 21.5 | 3198.2 ± 163.1 | 2442.8 ± 174.7 |
|  | **ChR2** | 84.8 ± 15.1 | 317.8 ± 26.4 | 182.6 ± 23.3 | 166.2 ± 22.4 | 3042.3 ± 183.6 | 2804.9 ± 196.0 |
| **Females** | **YFP** | 104.9 ± 10.1 | 256.8 ± 18.1 | 167.1 ± 13.9 | 108.7 ± 11.5 | 3908.9 ± 153.3 | 3858.1 ± 256.3 |
|  | **ChR2** | 93.3 ± 12.7 | 246.6 ± 22.7 | 166 ± 16.8 | 100.9 ± 17.6 | 4075.7 ± 231.0 | 3893.1 ± 153.5 |

**Table 2:** Exp. 1 Estrous Cycle Distribution. **Estrous cycle stages during optogenetic experiments.** Following assessments, vaginal cytology was used to approximate estrous cycle stage in cycling rats. RTPP: real-time place preference. n = 12/group.

| Group | Phase | RTPP | Social | Restraint | c-Fos |
| --- | --- | --- | --- | --- | --- |
| YFP | **Diestrus** | 33.3% | 25.0% | 33.3% | 25.0% |
|  | **Proestrus** | 16.7% | 33.3% | 33.3% | 58.3% |
|  | **Estrus** | 0.0% | 16.7% | 0.0% | 0.0% |
|  | **Metestrus** | 50.0% | 25.0% | 33.3% | 16.7% |
| ChR2 | **Diestrus** | 41.7% | 25.0% | 33.3% | 25.0% |
|  | **Proestrus** | 25.0% | 33.3% | 41.7% | 41.7% |
|  | **Estrus** | 0.0% | 16.7% | 8.3% | 8.3% |
|  | **Metestrus** | 33.3% | 25.0% | 16.7% | 25.0% |

**Table 3:** Exp. 2 Estrous Cycle Distribution. **Estrous cycle stages during TeNT experiment.** Following restraint, vaginal cytology was used to approximate estrous cycle stage in cycling rats. n = 8-10/group.

| Group | Phase | Restraint |
| --- | --- | --- |
| GFP | Diestrus | 0.0% |
|  | Proestrus | 50.0% |
|  | Estrus | 0.0% |
|  | Metestrus | 50.0% |
| TeNT | Diestrus | 0.0% |
|  | Proestrus | 50.0% |
|  | Estrus | 12.5% |
|  | Metestrus | 37.5% |

## Notes

### Competing Interest Statement

The authors have declared no competing interest.

